# Structure and conformational states of the bovine mitochondrial ATP synthase by cryo-EM

**DOI:** 10.1101/023770

**Authors:** Anna Zhou, Alexis Rohou, Daniel G. Schep, John V. Bason, Martin G. Montgomery, John E. Walker, Nikolaus Grigorieff, John L. Rubinstein

**Affiliations:** The Hospital for Sick Children Research Institute, Toronto, Canada; Department of Medical Biophysics, The University of Toronto, Canada; Howard Hughes Medical Institute, Janelia Research Campus, Ashburn, USA; MRC Mitochondrial Biology Unit, Cambridge, UK; Department of Biochemistry, The University of Toronto, Canada

## Abstract

Adenosine triphosphate (ATP), the chemical energy currency of biology, is synthesized in eukaryotic cells primarily by the mitochondrial ATP synthase. ATP synthases operate by a rotary catalytic mechanism where proton translocation through the membrane-inserted F_o_ region is coupled to ATP synthesis in the catalytic F_1_ region via rotation of a central rotor subcomplex. We report here single particle electron cryomicroscopy (cryo-EM) analysis of the bovine mitochondrial ATP synthase. Combining cryo-EM data with bioinformatic analysis allowed us to determine the fold of the a subunit, suggesting a proton translocation path through the F_o_ region that involves both the a and b subunits. 3D classification of images revealed seven distinct states of the enzyme that show different modes of bending and twisting in the intact ATP synthase. Rotational fluctuations of the c_8_-ring within the F_o_ region support a Brownian ratchet mechanism for proton-translocation driven rotation in ATP synthases.

## Introduction

In the mitochondria of eukaryotes, adenosine triphosphate (ATP) is produced by the ATP synthase, a ~600 kDa membrane protein complex composed of a soluble catalytic F_1_ region and a membrane-inserted F_o_ region. The ATP synthase is found in the inner membranes of mitochondria, with the F_1_ region in the mitochondrial matrix and the F_o_ region accessible from the inter-membrane space between the mitochondrial outer and inner membranes. In the mammalian enzyme the subunit composition is α_3_β_3_γδε for the F_1_ region with subunits a, e, f, g, A6L, DAPIT, a 6.8 kDa proteolipid, two membrane-inserted α-helices of subunit b, and the c_8_-ring forming the F_o_ region (1). The rotor subcomplex consists of subunits γ, δ, ε, and the c_8_-ring. In addition to the rotor, the F_1_ and F_o_ regions are linked by a peripheral stalk composed of subunits OSCP, d, F_6_, and the hydrophilic portion of subunit b. Approximately 85 % of the structure of the complex is known at high resolution from X-ray crystal structurs of constituent domains, which have been assembled into a mosaic structure within the constraints of a cryo-EM map at 18 Å resolution (1, 2).

The proton motive force, established by the electron transport chain during cellular respiration, drives protons across the F_o_ region through the interface between the a subunit and the c_8_-ring, inducing rotation of the rotor (3, 4). While the mechanism by which ATP synthesis and hydrolysis are coupled to rotation of the γ subunit is now understood well (1), it is still unresolved how rotation of the central rotor is coupled to proton translocation through the F_o_ region. The most popular model suggests that proton translocation occurs through two offset half channels near the a subunit/c subunit interface (5, 6). In this model, one half channel allows protons to move half-way across the lipid bilayer in order to protonate the conserved Glu58 residue of one of the c subunits. The other half channel allows deprotonation of an adjacent c subunit (7), setting up the necessary condition for a net rotation of the entire c-ring. Rotation does not occur directly from the protonating half channel to the deprotonating half channel, but in the opposite direction so that the protonated, and therefore uncharged, Glu residues traverse through the lipid environment before reaching the deprotonating half channel. The deprotonated Glu residue prevents the ring from rotating in the opposite direction, which would place the charged residue in the hydrophobic environment of the lipid bilayer. Rotation of the ring occurs due Brownian motion, making the enzyme a Brownian ratchet.

A recent cryo-EM map of the *Polytomella sp.* ATP synthase dimer showed two long and tilted α-helices from the a subunit in contact with the c_10_-ring of that species (8). This arrangement of α-helices from the a and c subunits was also seen in the *Saccharomyces cerevisiae* V-type ATPase (9). Cryo-EM of the *S. cerevisiae* V-ATPase demonstrated that images of rotary ATPases could be separated by 3D classification to reveal conformations of the complex that exist simultaneously in solution. In the work described here, we obtained and analyzed cryo-EM images of the bovine mitochondrial ATP synthase. 3D classification of the images resulted in seven distinct maps of the enzyme, each showing the complex in a different conformation. By averaging the density for the proton-translocating a subunit from the seven maps, we generated a map segment that shows α-helices clearly. Analysis of evolutionary covariance in the sequence of the a subunit (10) allowed the entire a subunit polypeptide to be traced through the density map. The resulting atomic model for the a subunit was fitted into the maps for the different rotational states, suggesting a path for protons through the enzyme and supporting the Brownian ratchet mechanism for the generation of rotation (5, 6), and thereby ATP synthesis, in the ATP synthase.

## Results

Specimens of ATP synthase were isolated from bovine heart mitochondria and prepared for cryo-EM as described previously (2, 11) (Fig.1 supplement 1). Initial 3D classification produced three classes, each of which appeared to show a ~120° rotation of the central rotor within the F_1_ region of the complex (Fig. 1A, blue arrows), similar to what was seen previously with the *S. cerevisiae* V-ATPase (9). Further classification of these three rotational states was able to separate state 1 into two sub-states, subsequently referred to as states 1a and 1b. State 2 could be divided into states 2a, 2b, and 2c, while state 3 could be separated into states 3a and 3b. Each of these 3D classes shows a different conformation of the enzyme (Fig. 1 supplement 2 and Movie 1). While the rotational states of the yeast V-ATPase were found to be populated unequally after 3D classification, bovine ATP synthase classes corresponding to different positions of the rotor had approximately equal populations. State 1 contained 43,039 particle images divided almost equally over its two sub-states, state 2 contained 48,053 particles images divided almost equally over its three sub-states, and state 3 contained 46,257 particle images divided almost equally over its two sub-states. The resolutions of the seven classes were between 6.4 and 7.4 Å (Fig. 1 supplement 3).

**Figure 1.**
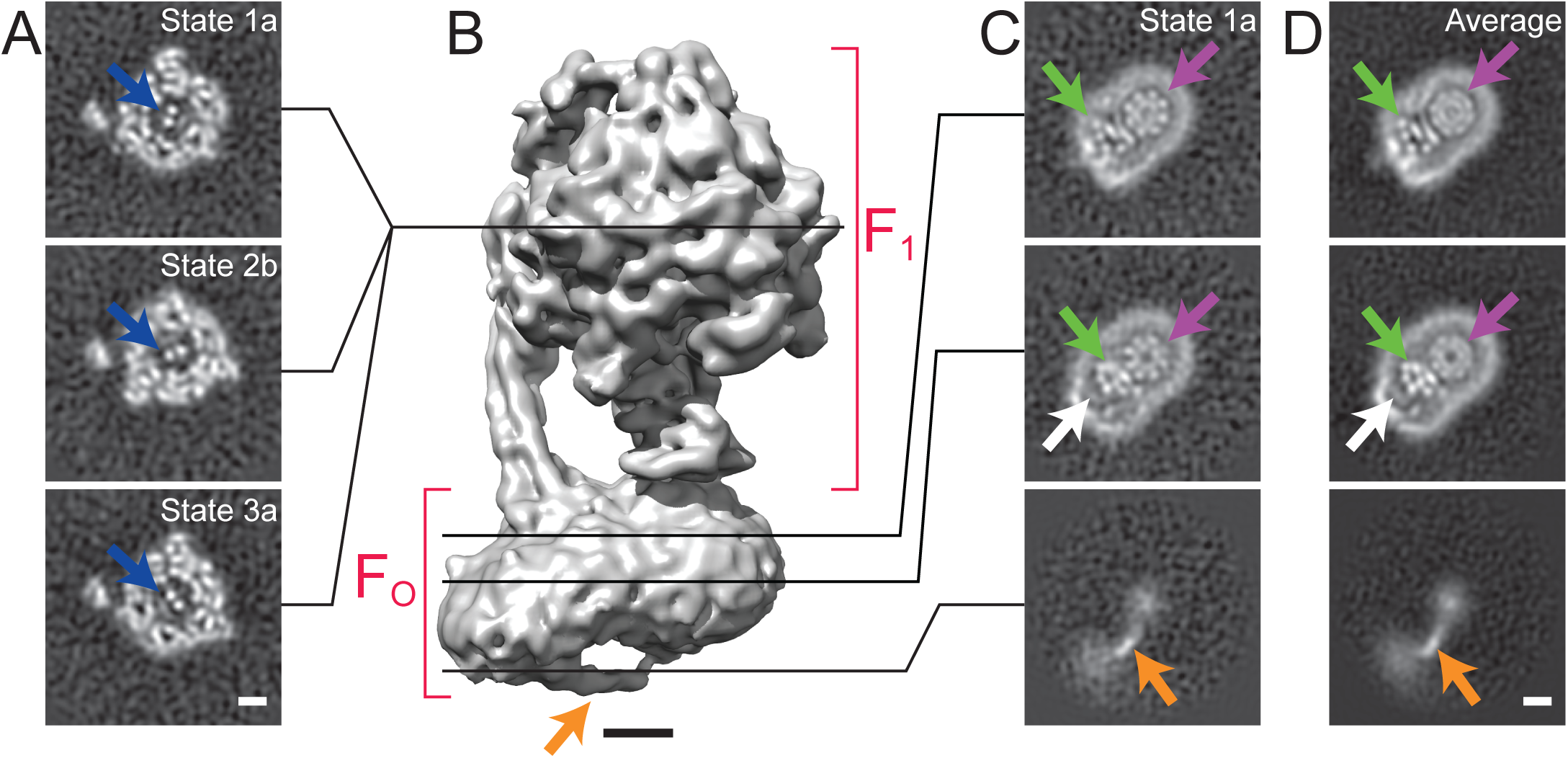
Cross-sections through maps. **A,** Cross-sections through the F_1_ regions of the different maps show that states 1, 2, and 3 are related by ~120° rotations of the y subunit within the α_3_β_3_ hexamer. **B**, Surface rendering of a map (State 1a) shows the bent F_o_ region with a tubular feature that extends from the rotor-distal portion to the c_8_-ring (orange arrow). **C**, Cross-sections through the F_o_ region shows α-helices from the a, b and A6L subunits (green arrows), a large low density region in the rotor-distal portion (white arrow), and the extension from the rotor-distal portion to the rotor (orange arrow). **D**, Averaging the F_o_ regions from the seven different maps shows all of the features mentioned above with an improved signal-to-noise ratio. Scale bars, 25 Å.

There is a distinct bend in the F_o_ region of the complex between the portion that is proximal to the c_8_-ring and the portion that is distal to the c_8_-ring (Fig. 1B and Fig. 1 supplement 2). This bent structure was seen previously in a lower-resolution cryo-EM map of the bovine mitochondrial ATP synthase (2). It is consistent with electron tomograms of ATP synthases in mitochondrial membranes (12, 13) and was also observed recently by electron tomography of membrane-reconstituted 2D crystals of the bovine enzyme (14). The e and g subunits are expected to reside in the portion of the F_o_ region distal to the c_8_-ring because cryo-EM maps of the mitochondrial ATP synthase from *S. cerevisiae,* where subunits e and g were removed by detergent, lacked this bent portion (2, 15). The f subunit is also thought to be associated with the e and g subunits (16). DAPIT and the 6.8 kDa proteolipid are not expected to be present in this preparation because the necessary lipids for maintaining their association were not added during purification (17, 18). While a detergent micelle can be seen around the entire F_o_ region, the portion of F_o_ distal to the c_8_-ring also contains a feature with unusually low density (Fig 1C, white arrows). The content of this low-density region is uncertain. The low density could be due to partial occupancy, flexibility, or disorder of a protein subunit. However, the low-density feature is bounded on one side by unusually sharp density from the detergent micelle, suggesting more order than in the rest of the micelle, and on the other side by the a subunit, which also appears well ordered. Instead, the low density could be due to bound material with low density, possibly lipid, that remains after purification of the enzyme.

### A novel feature in the F_o_ region

The F_o_ region of all seven maps also revealed a remarkable feature not resolved previously in cryo-EM maps of ATP synthases (2, 8, 15, 19). The feature appears to consist of an elongated membrane-embedded density, possibly an α-helix, that extends from the rotor-distal portion of F_o_ to the c_8_-ring. The orientation of this density would cause it to pass through the inter-membrane space of the mitochondrion (Fig. 1B and C, orange arrow). While not identified in the previous cryo-EM map of the enzyme at 18 Å resolution, the structure is consistent with a poorly-resolved ridge along the surface of F_o_ region seen in the earlier map (2). Because it extends from the bent end of the F_o_ region, this feature may correspond to the soluble part of the e subunit. Indeed, a similar structure was observed in single particle EM of negatively stained ATP synthase dimers from bovine heart mitochondria, and was proposed to be interacting e subunits (20). However, in the present structure the feature is not positioned to interact between dimers of the enzyme and its role in the complex remains unclear.

### Subunit a, b and A6L in the F_o_ region

In order to improve the signal-to-noise ratio for the F_o_ region of the complex, the membrane regions from the seven different maps were aligned and averaged. Averaging maps increases the signal-to-noise ratio where the structures are similar, but blurs regions where the maps differ. In principle, this method could also be applied to other map regions of the ATP synthase or other heterogeneous protein complexes by applying an appropriate transform before averaging. Averaging the F_o_ region provides a clear view of the portion of F_o_ region adjacent to the rotor, allowing the trans-membrane α-helices from the a, b, and A6L subunits to be identified reliably (Fig. 1C and D, green arrows, and Fig. 2). The c-ring has a lower density in the averaged membrane region than in the original maps, suggesting that its position differs between maps (Fig. 1C and D, ping arrows). The averaged density for the F_o_ region revealed the a subunit to have five membrane-inserted α-helices and an additional α-helix along the plane of the membrane surface (Fig. 2). Three additional trans-membrane α-helices are also apparent, presumably two from the b subunit (21) and one from the A6L subunit (22). The mammalian mitochondrial a subunit possesses the two highly tilted α-helices in contact with the c-ring that were seen previously for the *Polytomella sp.* F-type ATP synthase (8) and *S. cerevisiae* V-ATPase (9) (Fig. 2A).

**Figure 2.**
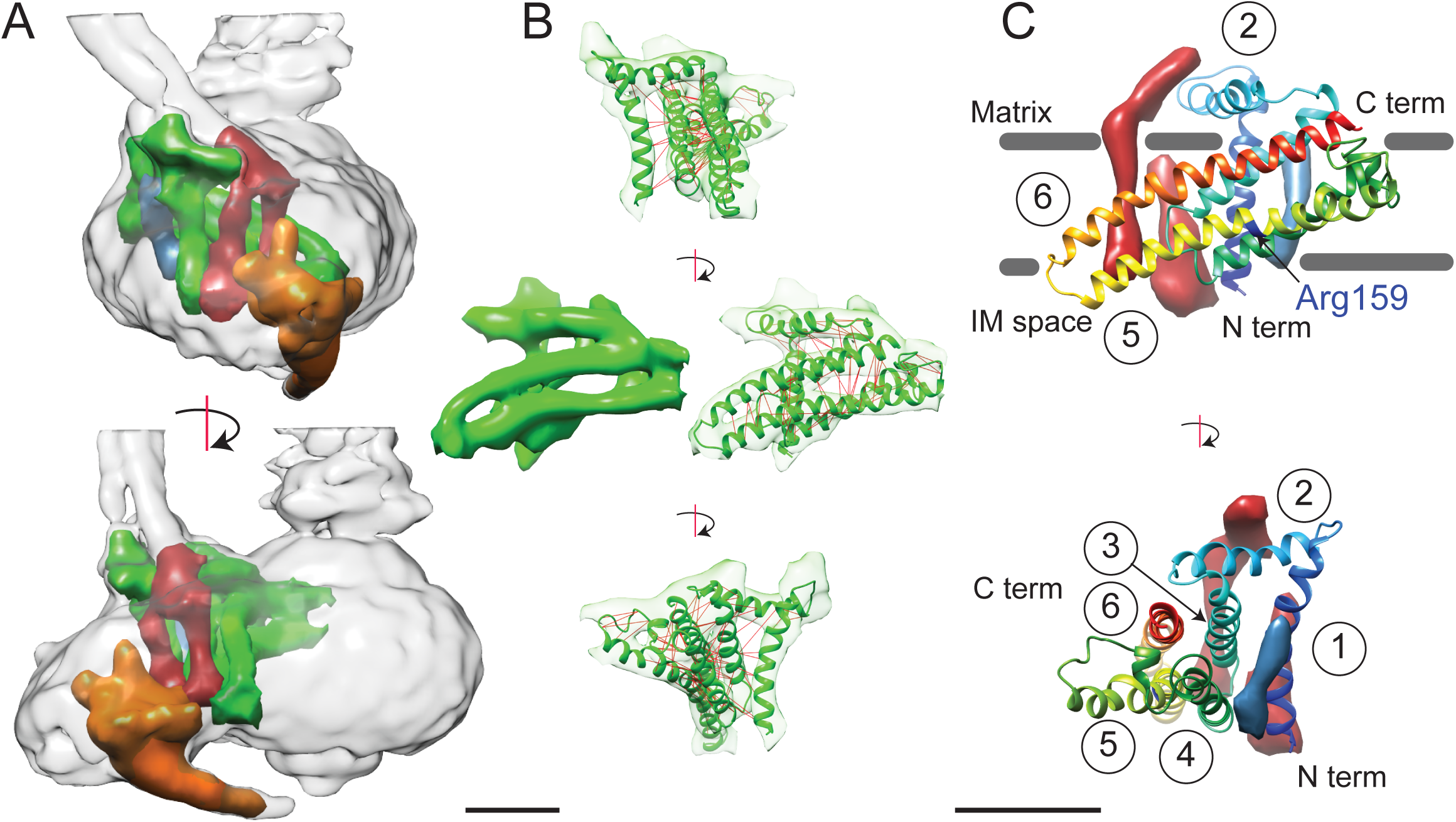
3D structure of the F_o_ region. **A**, In the F_o_ region of the complex, density was segmented for the a subunit (green), the b subunit (red-brown), the A6L subunit (blue), and the structure thought to arise from subunits e and g (orange). **B**, The a subunit sequence could be placed unambiguously into the cryo-EM density (green) by including constraints for residues predicted to be near to each other due to evolutionary covariance (red lines). **C**, The a subunit (coloured with a gradient from blue to red to denote directionality from the N to C terminus) possesses six α-helices, numbered 1–6. Trans-membrane α-helices from subunits b and A6L are shown as volumes (red-brown and blue, respectively). Five of the α-helices of subunit a are membrane-inserted while helix #2 runs along the matrix surface of the F_o_ region. The N terminus of the a subunit is on the inter-membrane space side of the subunit while the C terminus is on the matrix side. The highly conserved residue Arg159 is on the elongated and highly tilted α-helix #6. Scale bar, 25 Å.

A model for the a subunit was built into the cryo-EM density map using constraints from analysis of evolutionary covariance in sequences of the a subunit from different species. Analysis of covariance in evolutionarily related protein sequences can identify pairs of residues in a protein structure that are likely to interact physically with each other (10, 23–25). Spatial constraints from covariance analysis were sufficient not only to identify tentatively trans-membrane α-helices of the a subunit that are adjacent to each other, but also suggest which face each α-helix presents to the other α-helices (Fig. 2B and Movie 2, red lines). The constraints show patterns of interaction consistent with the predicted *α*-helical structure of the a subunit (Fig. 2 supplement 1A), as well as interactions between the a subunit and the outer α-helix of a c subunit in the c_8_-ring (Fig. 2 supplement 1B). As a result, we were able to trace, we believe unambiguously, the path of the a subunit polypeptide through the cryo-EM density map. The fit of the α-helices in the a subunit density was improved by molecular dynamics flexible fitting (*MDFF*) (26) and the 33 residue long connecting loop from residues 115 to 148 was built with *Rosetta* (27) (Fig. 2B and Movie 2). This connecting loop was built to be physically reasonable, but because its structure is not derived from experimental data it is not included in the discussion below. The final model places the α carbons of the residues in the co-varying pairs within 15 Å of each other in 94 % of the top 90 identified pairs, with an average C_α_ to C_α_ distance of 10.3 Å. The 6 % of constraints that are violated by the model are consistent with the false positive rate observed when testing covariance analysis approaches with proteins of known structure (28).

### Description of the a subunit structure

The mammalian a subunit appears to consist of six α-helices, with five α-helices that penetrate into the membrane-inserted portion of the enzyme (Fig. 2C). The N terminus of the subunit is in the inter-membrane space of the mitochondrion. The first α-helix extends vertically across the F_o_ region distal to the contact of the a subunit and c_8_-ring. The two trans-membrane α-helices of the b subunit are packed against one surface of helix #1 while the single trans-membrane α-helix from the A6L subunit is packed against its opposite surface. The second density region, interpreted as an α-helix of the a subunit, is not membrane-inserted and extends along the matrix surface of the F_o_ region connecting the membrane-inserted α-helix #1 with a membrane-inserted helical-hairpin composed of α-helices #3 and #4. This hairpin of the third and fourth α-helices does not appear to cross the F_o_ region fully, as seen previously in the *Polytomella sp.* ATP synthase (8). The final two trans-membrane helices are the two highly tilted α-helices seen previously with the *Polytomella sp.* ATP synthase and *S. cerevisiae* V-ATPase, with the C terminus of the a subunit on the matrix side of the F_o_ region. Within this structure, Arg159, which is essential and completely conserved, is found near the middle of the long tilted α-helix nearer the inter-membrane space side of the F_o_ region, different from its predicted position in the *Polytomella sp.* enzyme (8).

### Docking of atomic models into the cryo-EM maps

To analyze the different enzyme conformations detected by 3D classification, the maps were segmented and available crystal structures for the F_1_:IF_1_ complex (29), F_1_ peripheral stalk complex (30), peripheral stalk alone (31), and F_1_-c_8_ complex (32) were combined into each of the maps by *MDFF* (26). Residues for the b subunit were extended from the N terminus of the b subunit crystal structure into the membrane region based on trans-membrane α-helix prediction. While *MDFF* with maps in this resolution range cannot be used to determine the locations or conformations of amino acid side chains, loops, or random-coil segments of models, it can show the positioning of α-helices in the structures. Figures 3A and B compare the fitting for state 1a (Fig. 3A) and state 1b (Fig. 3B), illustrating the accuracy with which α-helical segments could be resolved in the maps of different sub-states. The atomic model alone for state 1a is shown in Figure 3C, with the c_8_-ring removed for clarity in Figure 3D. Transitions between the different states were illustrated by linear interpolation (Movie 3). As seen previously for the *S. cerevisiae* V-ATPase, almost all of the subunits in the enzyme undergo conformational changes on transition between states (9). Because there were two sub-states identified for states 1 and 3 there is only a single sub-state to sub-state transition for these two states. In comparison, three different sub-states were identified for state 2 and consequently there are three sub-state to sub-state transitions that are possible for this state. All of the sub-state to sub-state transitions include a slight rotation of the c_8_-ring against the a subunit. It is possible that this movement is due to partial disruption of the subunit a/c_8_-ring interface. However, the structural differences within the F_o_ regions of different classes are significantly smaller than the structural differences seen elsewhere in the enzyme, suggesting that these changes do not originate from disruption within the membrane region of the complex and instead reflect flexibility in the enzyme.

**Figure 3.**
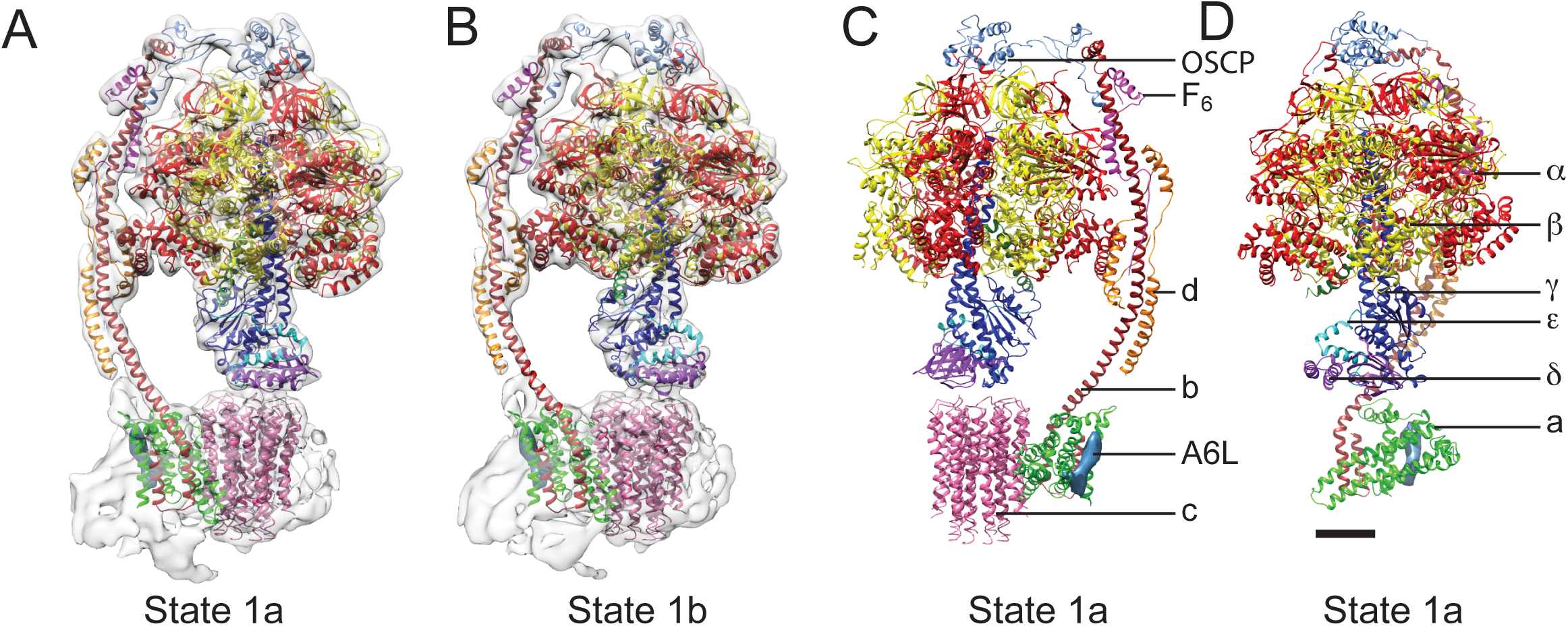
Docking of atomic models into the cryo-EM maps. Fitting of all available atomic models into the density map is shown for state 1a (**A**) and state 1b (**B**). State 1a is also shown in a different orientation and without the density map (**C**) and with the c_8_-ring removed for clarity (**D**). The apparent gap between the c_8_-ring and γ and δ subunits is filled with amino acid side chains and is the same as was seen in the crystal structure of the F_1_-c_8_ complex (32). Scale bar, 25 Å.

The largest change between the sub-states of each state can be approximated by rotations of the α_3_β_3_ hexamer relative to the rest of the complex by angles ranging from 10 to 16°. A comparison of the maps for the different sub-states and the axes of these rotations are shown in Figure 3 supplement 1. The resulting conformational changes can be seen most clearly in Movies 4 and 5. The state 1a to 1b transition reveals a bending of the peripheral stalk towards the top of the F_1_ region near the OSCP and F_6_ subunits (Movies 4 and 5, panel A). In comparison, the state 3a to 3b transition reveals bending of the peripheral stalk near where the b subunit enters the membrane (Movies 4 and 5, panel D). Transitions between the three sub-states of state 2 show both motions: the transition between 2a and 2b shows mostly bending of the peripheral stalk near OSCP and F_6_ subunits while the 2b and 2c transition shows mostly bending near the membrane inserted portion of the peripheral stalk (Movies 4 and 5, panels B and C, respectively). The transition from state 2a to 2c shows a combined bending at both of these positions. It is most likely that the different modes of bending exist in all of the states and further classification of larger datasets would be expected to reveal these complex motions. The different sub-states do not appear to have a specific sequence or represent specific intermediates in the rotation sequence. Instead, the differences in conformation between sub-states when taken together illustrate the flexibility of the enzyme, a property that has been linked to its rapid rate of enzymatic activity (9, 33). The functional significance of the sub-states may also be determined by the orientation of the c_8_-ring with respect to the a subunit, as discussed below.

### Discussion

Predicting the path of protons through membrane protein complexes has proven difficult, even in cases where high-resolution atomic models including bound water molecules are available from X-ray crystallography (34). Nonetheless, features in the structure of the bovine mitochondrial F_o_ region suggest a possible path for proton translocation similar to a model put forward based on the structure of the *Polytomella sp.* ATP synthase (8). The arrangement of α-helices in the F_o_ region is remarkably similar to the arrangement of a-helices in the V_o_ region of the yeast V-ATPase (9), even though the V-ATPase a subunit has eight α-helices and little detectable sequence similarity with the F-type ATP synthase a subunit. The conserved general architecture of the membrane-inserted regions in F-type ATP synthases and V-type ATPases suggests that the observed arrangement of α-helices is functionally important and likely involved in proton translocation (Fig. 4A and B). The matrix half channel of the ATP synthase is likely to be formed by the cavity between the c_8_-ring and the matrix ends of tilted α-helices #5 and #6 of the a subunit. The luminal half channel in the V-ATPase is probably formed entirely from α-helices from the a subunit, whereas the corresponding inter-membrane space half channel in the ATP synthase is likely composed of the inter-membrane space ends of α-helices #5 and #6 and one or both of the two trans-membrane α-helices of the b subunit. Defining the exact placement of half channels will likely require higher-resolution maps from cryo-EM or X-ray crystallography that reveal amino acid side chain density and bound water molecules.

**Figure 4.**
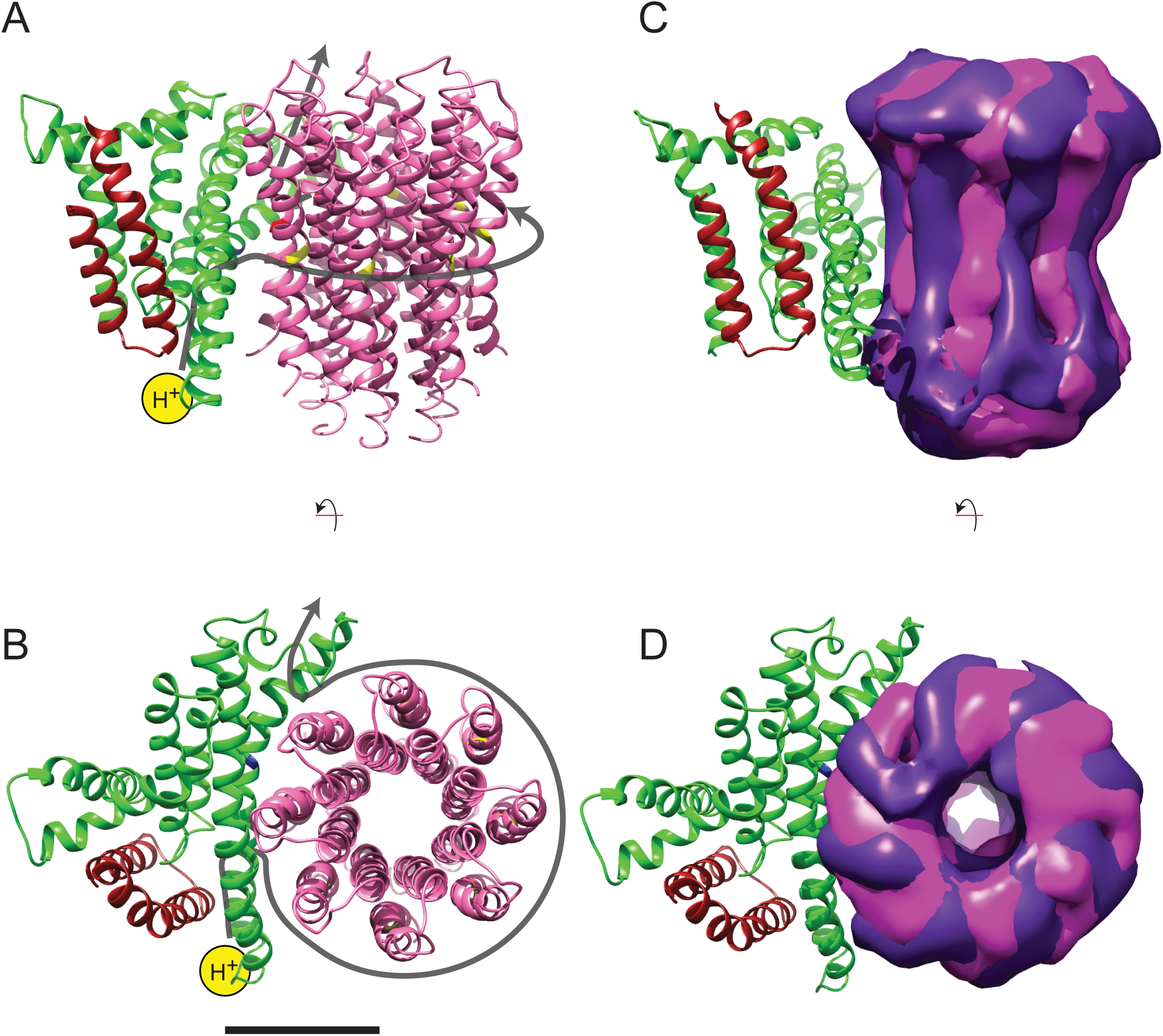
Model for proton translocation. **A**, The a subunit, along with the membrane-intrinsic α-helices of the b subunit, form two clusters that could be the half channels needed for trans-membrane proton translocation. **B**, The map segment corresponding to the c_8_-ring is shown for state 2a (pink) and state 2c (purple). The difference in rotational position of the ring is consistent with the Brownian fluctuations predicted for the generation of a net rotation. Scale bar, 25 Å.

In addition to bending and twisting of the peripheral stalk and central rotor of the enzyme, the differences between the sub-states of each state show variability in the rotational position of the c_8_-ring in relation to the a subunit (Fig. 4C and D), even in the nucleotide-depleted conditions in which cryo-EM grids were frozen for this analysis. This lack of a rigid interaction between the c_8_-ring and a subunit is consistent with the Brownian ratchet model of proton translocation (5). In the Brownian ratchet model, the rotational position of the ring fluctuates due to Brownian motion, but cannot turn to place the Glu58 residue of a c subunit into the hydrophobic environment of the lipid bilayer until the Glu58 is protonated by the inter-membrane space half-channel. Therefore, with this model, the different sub-states would correspond to energetically equivalent or nearly-equivalent conformations that occur due to Brownian motion. Movie 6 illustrates the extent of rotational oscillation predicted from the transition between states 2a and 2c. It is most likely that this oscillation occurs as each c subunit passes the interface with the a subunit, with 8/3 c subunits on average contributing to the synthesis of one ATP molecule. The rotational flexibility of the c_8_-ring that exists even when the γ subunit is locked within the α_3_β_3_ hexamer suggests that flexing and bending of the components of the ATP synthase smooths the coupling of the 8-step rotation of the c_8_-ring with the 3-step rotation of the F_1_ region. This model suggests that the observed flexibility in the enzyme, which apparently complicates determination of atomic resolution structures directly from cryo-EM data, is also essential to the mechanism of ATP synthesis.

## Methods

### Protein purification and electron microscopy

Bovine mitochondrial ATP synthase was purified as described previously (11) and cryo-EM specimen grids were prepared as described previously, except that glycerol was removed from specimens prior to grid freezing with a 7 kDa molecular weight cutoff Zeba Spin centrifuged desalting column (Thermo Scientific) and nano-fabricated grids with 500 nm holes were used (35). After optimization of grid freezing conditions, micrographs were recorded from three grids on a Titan Krios microscope (FEI) operated at 300 kV with parallel illumination of a 2.5 μm diameter area of the specimen and an electron fluency of 3 el^−^/Å^2^ /s. A 70 μm objective aperture was employed with a nominal magnification of 18,000× onto a K2 Summit direct detector device (Gatan Inc.) operated in super-resolution mode with a 1.64 Å physical pixel and 0.82 Å super-resolution pixel. With no specimen present, the rate of exposure of the detector was 8 el^-^/pixel/s. Exposure-fractionated movies of 20.1 s were recorded as stacks of 67 frames, so that selected specimen areas were exposed with a total of 60.3 el^−^/Å^2^. Data collection was automated with *SerialEM* (36).

### Image processing

Magnification anisotropy (37) under the conditions described above was measured previously from images of a standard cross-grating specimen with the program *mag_distortion_estimate* (38). The linear scaling parameters were 0.986 and 1.013, the azimuth of the distortion was 134.0°, and the program *mag_distortion_correct* was used to correct for this distortion in each dose-fractionated frame. The frames were then down-sampled to a pixel size of 1.64 Å by Fourier-space cropping and aligned with each other with the program *Unblur* (39). Defocus parameters were estimated from aligned sums of frames using *CTFF1ND4* (40). Particle images were selected in *Relion* and subjected to 2D classification (41, 42), yielding a set of 195,233 single particle coordinates selected from 5,825 movies. Local beam-induced motion was corrected for each particle with the program *alignparts_lmbfgs* (43). Aligned dose-fractionated particle images were filtered and summed to optimize the signal-to-noise ratio at all spatial frequencies (39, 43, 44), giving a set of particle images that were 256×256 pixels. These images were down-sampled to 128×128 pixels (pixel size of 3.28 Å) for determining particle orientations. Initial single-particle alignment parameter values were obtained by 5 rounds of iterative grid search and reconstruction in *FREALIGN*'s mode 3 (45), using the earlier published map of the enzyme as an initial reference (2). *FREALIGN*'s likelihood-based classification algorithm (46) was then used to classify particles images into several maps, alternating between refinement of orientation parameters every 3^rd^ or 4^th^ iteration and class occupancy during other iterations. The final classification yielded 12 classes, of which 7 gave interpretable 3D maps. Only spatial frequencies up to 1/10 Å^-1^ were used during refinement to avoid fitting noise to high-resolution features in maps. All seven 3D maps had Fourier shell correlation values greater than 0.8 at this frequency.

### Map analysis and model building

Segmentation of 3D maps was performed with *UCSF Chimera* (47, 48) and atomic structures were fit flexibly into 3D maps using *NAMD* with Molecular Dynamics Flexible Fitting *(MDFF)* (26). The F_o_ regions from the seven different 3D maps were aligned and averaged in real space with *UCSF Chimera* and *Situs* (49). Co-varying pairs of residues were detected in the full bovine mitochondrial ATP synthase a subunit sequence (NCBI reference YP_209210.1) with the program *EVcouplings* (24) using a pseudo-likelihood maximization approach and the top 90 connections were considered in the analysis. The protein was not assumed to have trans-membrane α-helices and the job was run as a quick launch with all other parameters at default settings. Evolutionary couplings between the a subunit and ATP synthase c subunit were detected with *GREMLIN* (25) with an E-value threshold for multiple sequence alignments (MSAs) of 1×10^-10^ and *Jackhmmer* was used to produce MSAs over 8 iterations.

To build a model of the a subunit, six straight α-helices (*ϕ*=-57° and *ψ*=-47°) were built in *UCSF Chimera*. These α-helices were fit manually in the map of the average F_o_ region in the only orientations that satisfied constraints from evolutionary covariance analysis. For illustration, but not interpretation, loops connecting these helices were also included in the model. Randomly structured connecting loops between the α-helices were built in *Modeller* (50) within *UCSF Chimera* and fitted into the density with *MDFF* with a low density scaling factor (gscale=0.3) over 200,000 steps (200 ps). Bond lengths and angles were then idealized with *Rosetta* (idealize_jd2 command) and the loop between residues 115 and 148 rebuilt in *Rosetta* (loopmodel command) using the quick_ccd method of remodelling (51). Each output structure included an all-atom relaxation in the density map with a score weight of 0.1. The lowest energy models of 100 models was selected and angles were idealized and the structure energy-minimized with *UCSF Chimera.* Loops beside the one from residues 115 and 148 were too short for this process to be useful. The b subunit crystal structure was extended into the F_o_ region of the map based on trans-membrane α-helix prediction from MEMSAT-SVM (52).

## Acknowledgements

We thank Richard Henderson and Voula Kanelis for a critical reading of this manuscript. A preprint of this manuscript was first deposited on bioRxiv.org (http://dx.doi.org/10.1101/023770) on August 11, 2015. This work was supported by operating grant MOP 81294 from the Canadian Institutes of Health Research (JLR) and Medical Research Council grant U105663150 (JW). AZ was supported by a postgraduate scholarship from the Canadian Institutes of Health Research, an award from The Hospital for Sick Children, and a U of T excellence award. DGS was supported by a postgraduate scholarship from the Natural Sciences and Engineering Research Council and an award from The Hospital for Sick Children. JLR holds the Canada Research Chair in Electron Cryomicroscopy.

## Author contributions

JW, NG, and JLR initiated the projected. JVB and MGM purified the protein. AZ prepared specimens, performed automated particle picking, segmented maps, docked atomic models into maps, and analyzed conformational states. AR imaged the specimens and calculated 3D maps. DGS built the atomic model of the a subunit using information from evolutionary covariance and *Rosetta.* JLR, NG, and JW supervised the research.

## Author information

Cryo-EM maps have been deposited in the Electron Microscopy Data Bank with accession numbers EMDB-3164 to 3170. Atomic models have been deposited in the Protein Data Bank with accession numbers 5ARA, 5ARE, 5ARH, 5ARI, 5FIJ, 5FIK, and 5FIL. The authors declare no competing interests, financial or otherwise.

**Figure Captions**

**Figure 1 supplement 1.**
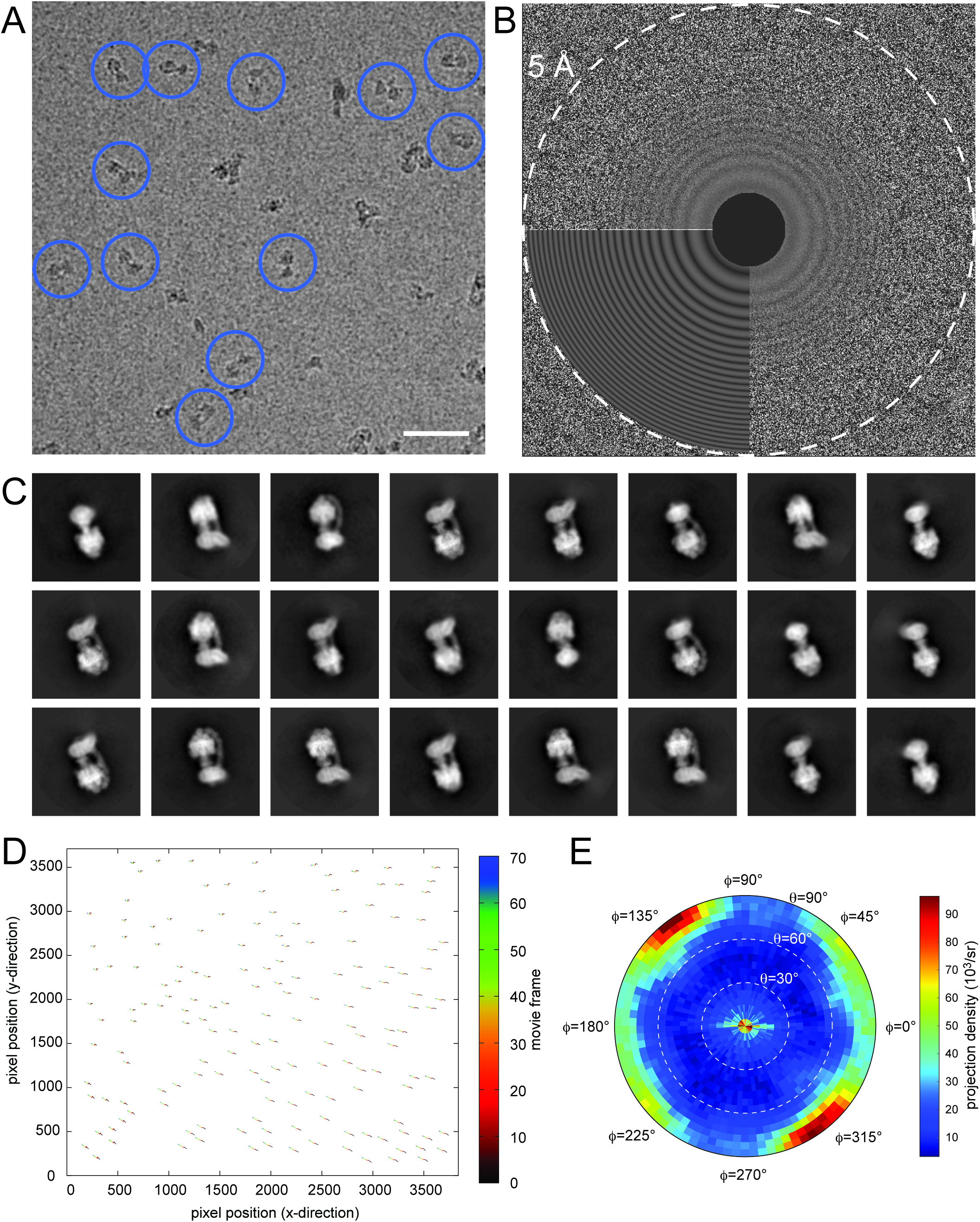
Electron microscopy and model construction. **A**, A sample micrograph with representative ATP synthase particles circled in blue. Scale bar, 500 Å. **B**, A sample micrograph power spectrum with the modeled power spectrum from *CTFF1ND4* in the lower left quadrant. **C**, Representative averages from 2D classes selected for 3D analysis. **D**, An example of individual particle trajectories (exaggerated by a factor of 5) from *alignparts_lmbfgs* used for local drift correction. **E**, The distribution of Euler angles assigned to particles going into the 3D maps shows preferred orientations but contains sufficient particle views to produce maps with isotropic resolution.

**Figure 1 supplement 2.**
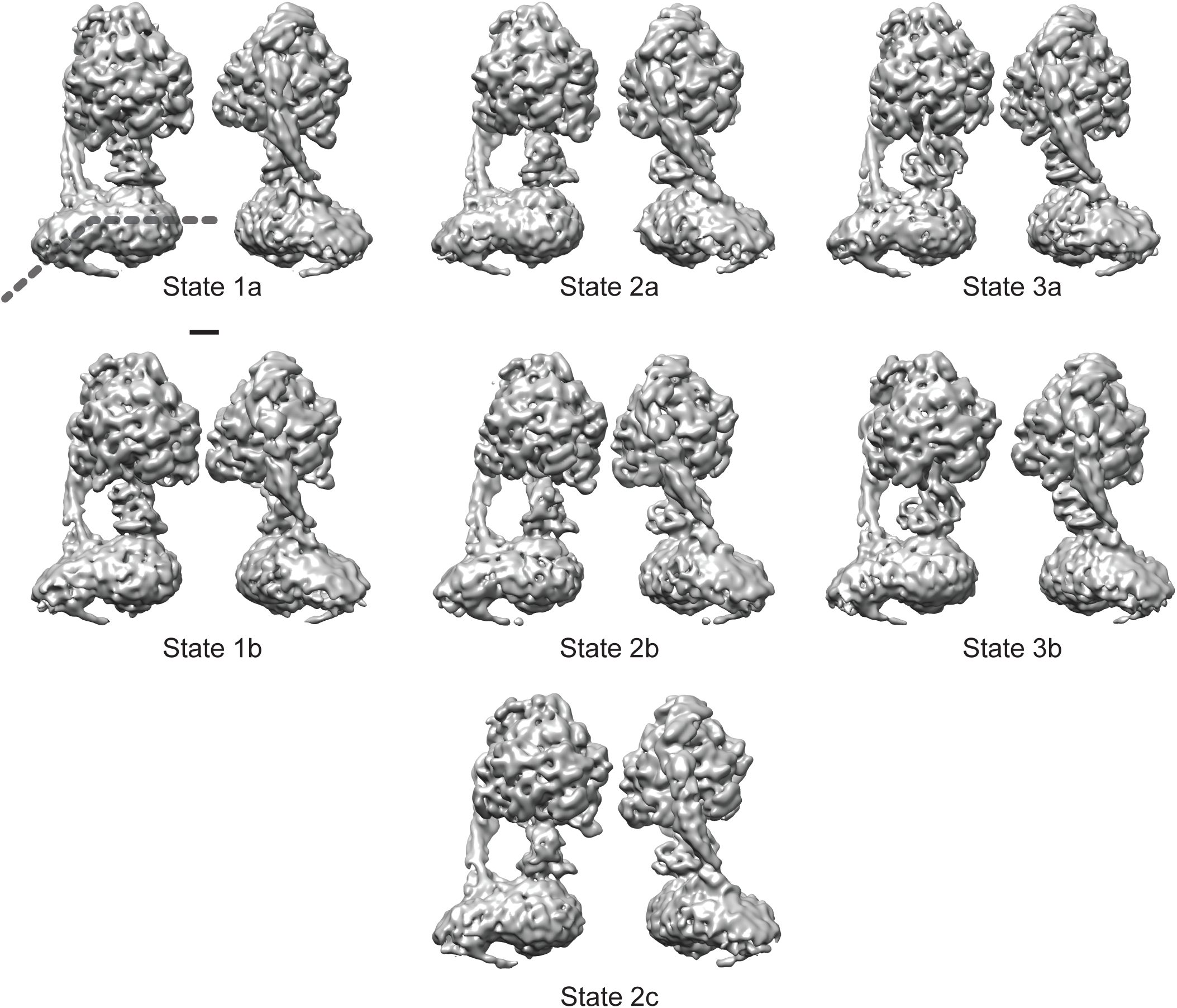
The seven observed states of the bovine mitochondrial ATP synthase. Two views are shown for each of the seven conformations identified for the enzyme. All of the known structural features and the newly observed protuberance from the rotor-distal portion of the F_o_ region are seen in each map. The bend in the F_o_ region is indicated by the dashed line in the top left map. Scale bar, 25 Å.

**Figure 1 supplement 3.**
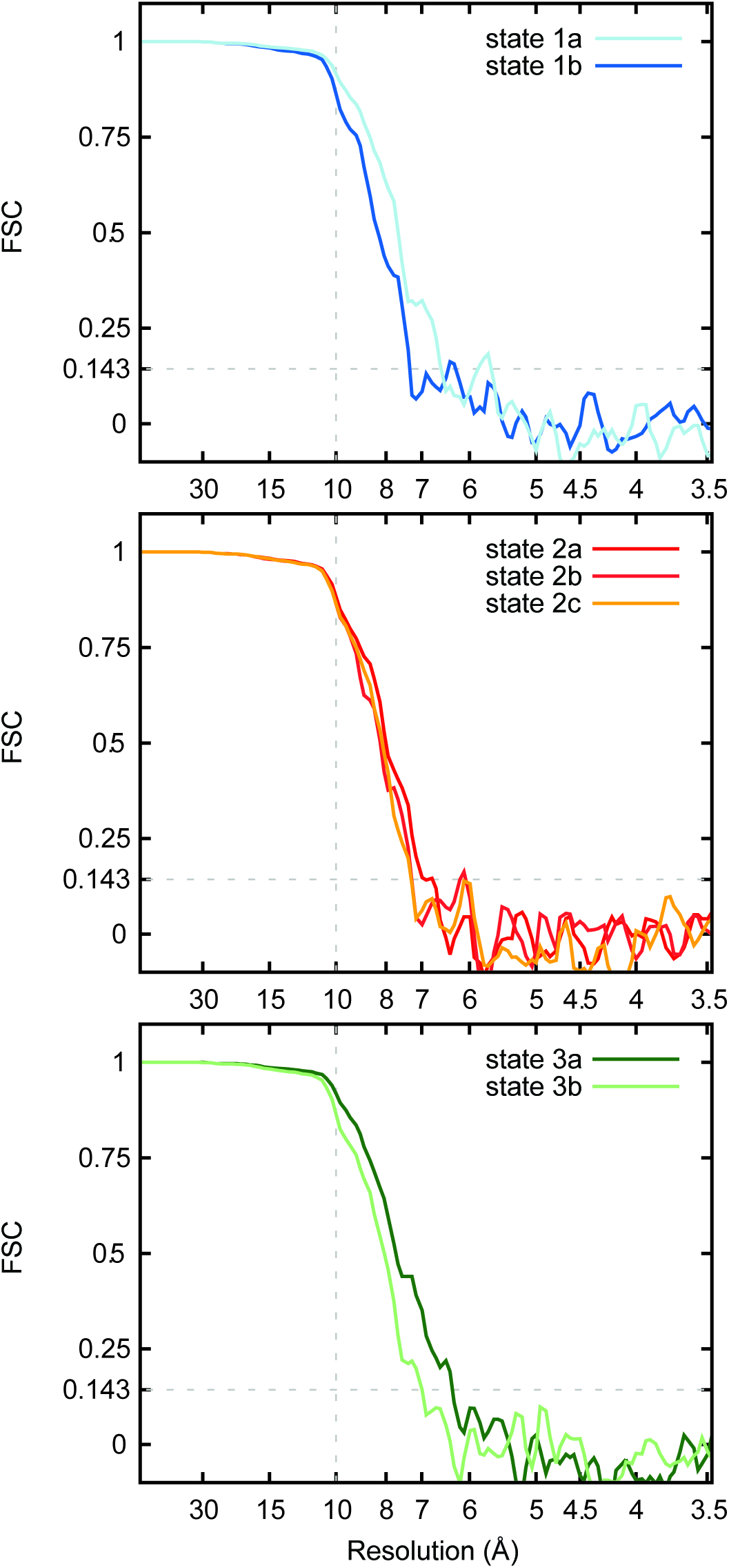
Fourier shell correlation curves for the seven maps. FSC curves are shown for state 1a and 1b (**A**), state 2a, 2b, and 2c (**B**), and state 3a and 3b (**C**).

**Figure 2 supplement 1.**
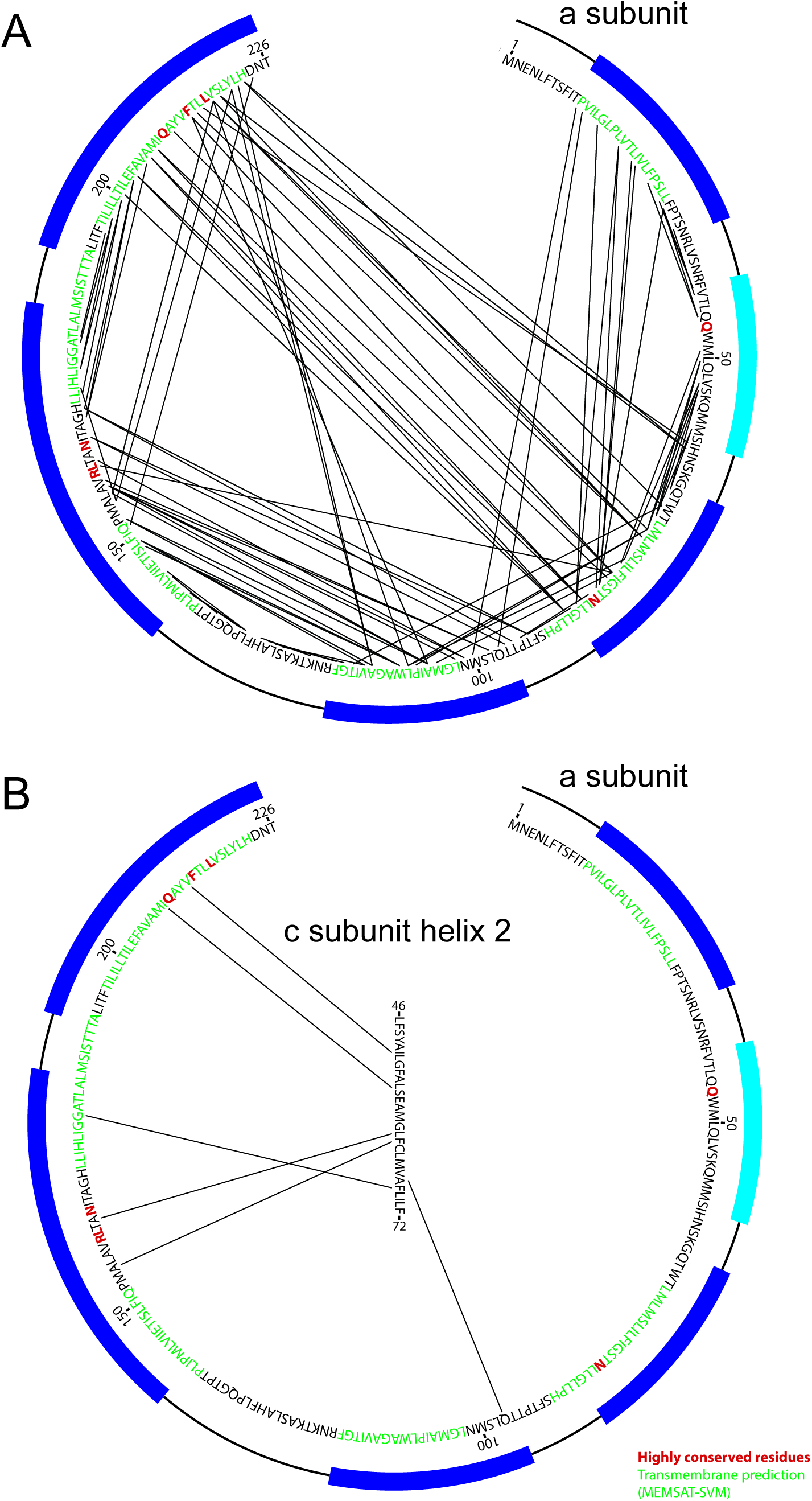
Analysis of evolutionary covariance of residues. **A**, The top 90 predicted couplings between residues of the a subunit are indicated, along with trans-membrane helices predicted by the MEMSAT-SVM algorithm (52), shown in green, and highly conserved residues, shown in red. Residues modeled as membrane-inserted a-helices based on the cryo-EM density are indicated with dark blue rectangles outside of the sequence and residues modeled as a soluble α-helix based on the cryo-EM density is indicated with a light blue rectangle. **B**, The top six predicted couplings between residues of the a subunit and residues on the outer surface of the c-ring are indicated.

**Figure 3 supplement 1.**
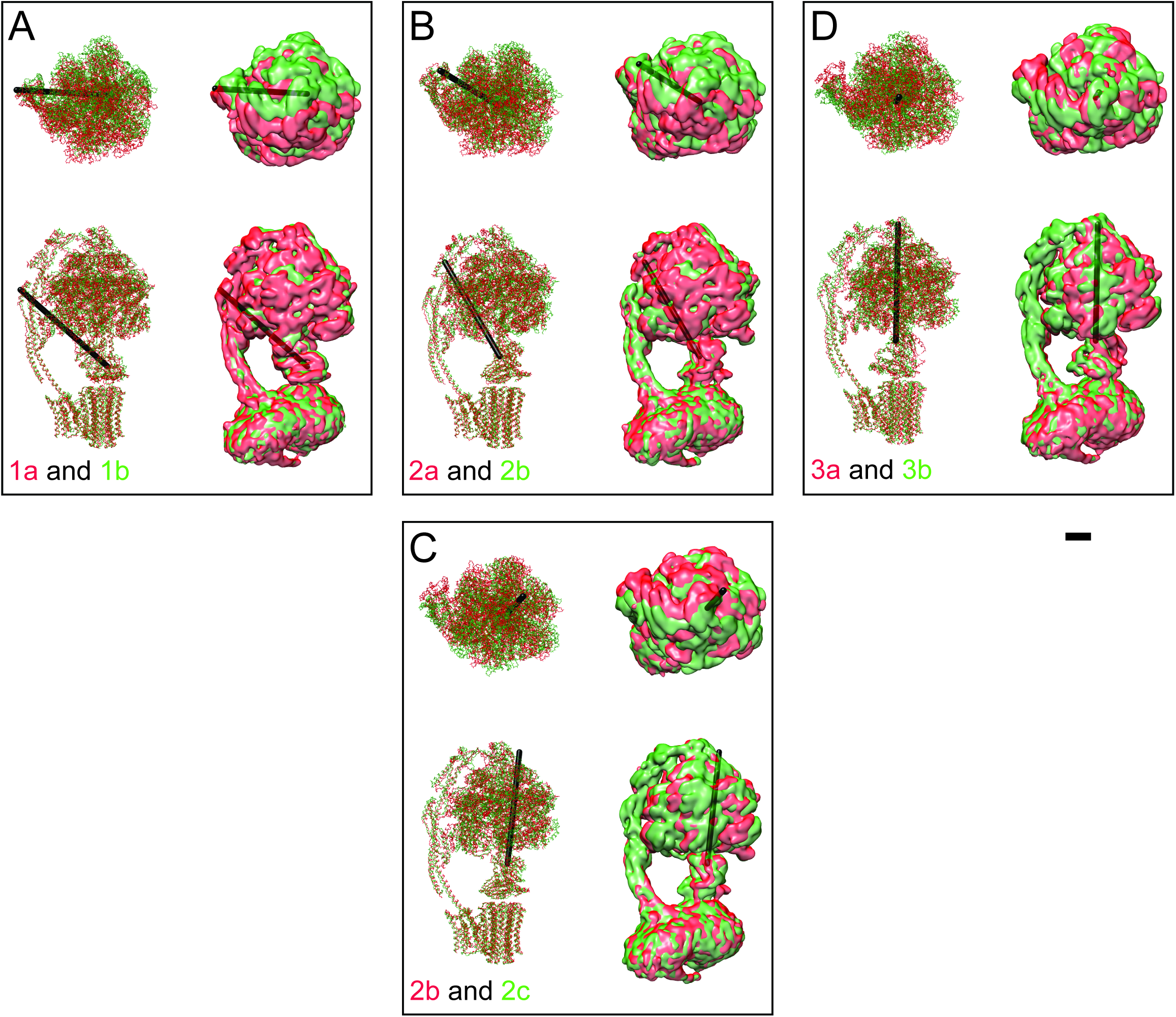
Differences between sub-states. The differences between sub-states can be seen by overlaying maps and models for state 1a (red) and 1b (green) (**A**), 2a (red) and 2b (green) (**B**), 2b (red) and 2c (green) (**C**), and 3a (red) and 3b (green) (**D**). These differences can be represented approximately as a rigid body rotation of the α_3_β_3_ hexamer by 10° (state 1a to 1b), 11° (state 2a to 2b), 12° (state 2b to 1c), and 16° (state 3a to 3b). The axes of these rotations are shown as black rods. This movement is most easily seen in Movies 4 and 5. Scale bar, 25 Å.

---

**Movie 1. Slices through the seven cryo-EM maps.** Cross-sections are shown moving from the F_1_ region towards the F_o_ region for states 1a, 1b, 2a, 2b, 2c, 3a, and 3b.

**Movie 2. Fold of the a subunit.** The density corresponding to the a subunit (green) and membrane-inserted portion of the b subunit (red-brown) and A6L subunit (blue) are shown, in addition to a ribbon diagram for the a subunit (green) and the top 90 constraints from analysis of covarying residues in the a subunit sequence (red lines). 6 % of the constraints could not be satisfied, which is consistent with the false positive rate from known structures (28). Scale bar, 25 Å.

**Movie 3. Conformation changes during the rotary cycle.** Linear interpolation is shown between one of each of the main states identified by 3D classification (state 1a, 2a, and 3a) showing the large conformation changes that occur during rotation. Scale bar, 25 Å. [*please view movie as loop*]

**Movie 4. Conformational differences between the different ATP synthase maps.** Differences between conformations detected by 3D classification are illustrated by linear interpolation between state 1a and 1b, showing bending of the peripheral stalk near the OSCP and F_6_ subunits (**A**), state 2a and 2b showing a similar bending of the peripheral stalk near the OSCP and F_6_ subunits (**B**), state 2b and 2c showing bending of the peripheral stalk near the membrane region (**C**), and state 3a and 3b showing a similar bending of the peripheral stalk near the membrane region (**D**). Scale bar, 25 Å. *[please view movie as loop*]

**Movie 5. Conformational differences between the different ATP synthase maps.** The same interpolations as shown in movie 4, except viewed from the F_1_ region towards the F_o_ region. Scale bar, 25 Å. *[please view movie as loop]*

**Movie 6. Brownian ratcheting.** The different rotational positions of the c_8_-ring between the sub-states are illustrated by interpolating between the positions of states 2a and 2c. The conserved residue Arg159 is shown as a blue sphere. The movement is consistent with the Brownian ratcheting predicted during proton translocation in rotary ATPases. Glu58 residues are shown moving from the proton-locked conformation (53) to an open conformation (54, 55) when they are close to the conserved Arg159 residue. Scale bar, 25 Å. [*please view movie as loop*]

